# Genetic and epigenetic divergence among *Arabidopsis thaliana* Col-0 laboratory lineages since the 1950s

**DOI:** 10.64898/2026.08.05.743031

**Authors:** Alexandra M. Tadros, Zhilin Zhang, Nan Yao, Rebecca A. Mosher, Frank Johannes, Robert J. Schmitz

## Abstract

The model plant *Arabidopsis thaliana* has been a workhorse of plant biology since the 1950s, when the Columbia-0 (Col-0) accession was widely adopted by the research community as a reference genotype. Since then, Col-0 seeds have been shared among laboratories worldwide and propagated independently, creating a network of related laboratory lineages. Here, we sequenced the genomes and DNA methylomes of 78 Col-0 strains from laboratories and stock centers around the world to assess the genetic and epigenetic variation accumulated during this period and to reconstruct their propagation history from *de novo* variants. After quality filtering, we identified 1,415 single-nucleotide polymorphisms and 169,677 epimutations across 75 Col-0 lines and used these variants to generate SNP- and epimutation-based phylogenies. The two phylogenies showed highly concordant topologies, but the epimutation-based tree provided greater resolution of relationships among closely related lineages. Divergence-time estimates suggest that many lines have been propagated for approximately one generation per year over several decades, resulting in pairwise divergences of up to approximately 100 generations. We further identified 170 nonsynonymous substitutions and more than 200 candidate epialleles, some of which were associated with gene-expression differences among Col-0 clades and may therefore have functional consequences. Together, these findings show that Col-0 is best viewed not as a single invariant reference genotype, but as a collection of related laboratory lineages that have accumulated genetic, epigenetic, and transcriptional divergence through time, with potential consequences for the reproducibility of experiments across laboratories.

**Significance Statement:** The *Arabidopsis* Col-0 reference accession is often treated as a single, invariant genotype, yet independently maintained laboratory stocks have accumulated substantial genetic, epigenetic, and transcriptional divergence since the 1950s. Some lineages are separated by more than 100 generations of independent propagation. Mutations and CG epimutations reconstruct nearly identical lineage histories, demonstrating that heritable DNA methylation changes provide a robust record of recent laboratory evolution.

## Introduction

*Arabidopsis thaliana* was first studied in a genetic context in Germany in 1907, when Friedrich Laibach correctly identified its five chromosome pairs (1). His interest in *A. thaliana* resumed several decades later (2), and in 1943 he described several features that made the species well suited to genetic research, including its small size, ease of cultivation, high progeny number and short generation time (3). Over the course of his career, Laibach and his colleagues collected *A. thaliana* seeds from across Europe. He maintained a well-organized collection of more than 150 accessions, which became the basis of the first Arabidopsis seed bank associated with the Arabidopsis Information Service (4, 5).

In the 1950s, the Hungarian scientist George Rédei became interested in *A. thaliana* and received seeds from Laibach, which he brought to the University of Missouri (USA) where he established his laboratory in 1956 (6). Laibach had provided seeds from four accessions, including Landsberg. After irradiating some of the Landsberg seeds, Rédei found that the material contained more than one genotype. One mutant with an erect growth habit became the foundation of the Landsberg *erecta* line. From non-mutagenized material, Rédei established the Columbia wild-type line from which Columbia-0 (Col-0) originated (6).

Interest in *A. thaliana* increased during the 1960s and 1970s, but its use expanded particularly rapidly following the development of *Agrobacterium tumefaciens*-mediated transformation in 1986 (7). As the Arabidopsis research community grew, Columbia material was exchanged among laboratories and independently propagated. Multiple seed stock centers were established in 1991, including the Arabidopsis Biological Resource Center (ABRC) and the Nottingham Arabidopsis Stock Centre (NASC), which maintain collections and provide an organized system for researchers to deposit and obtain seeds (8). Both stock centers and individual laboratories maintain their seed supplies through repeated propagation. Contemporary Col-0 stocks therefore form a network of related laboratory lineages connected through historical seed exchange but separated by decades of independent maintenance.

Repeated propagation provides opportunities for genetic mutations and heritable epigenetic changes, or epimutations, to arise and accumulate (9–15). Although many such changes are likely to be functionally neutral, others may alter protein function, gene regulation or phenotype. Thus, independently maintained Col-0 stocks may differ genetically, epigenetically and functionally despite being treated as the same reference genotype. Previous work identified genetic differences among individual Col-0 laboratory stocks and showed that these can contribute to inconsistent phenotypic responses across laboratories (16). However, the broader extent and structure of genetic and epigenetic variation across the worldwide collection of Col-0 stocks remains poorly understood. It also remains unclear whether this accumulated molecular divergence is accompanied by transcriptional differences or whether potentially functional variants are restricted to particular lineages and loci.

Here, we sequenced the genomes and DNA methylomes of 78 Col-0 seed stocks obtained from laboratories and stock centers around the world and characterized the genetic, epigenetic and transcriptional variation that had accumulated during their independent propagation. We used SNPs and single-site CG epimutations to examine relationships and propagation histories among the stocks, quantified coding mutations and regional methylation variants representing candidate epialleles, and tested whether accumulated genetic and epigenetic differences were associated with transcriptional variation. Together, these analyses assessed whether Col-0 could still be regarded as a single invariant reference genotype and highlighted the implications of long-term propagation and seed exchange for experimental reproducibility. More broadly, similar questions arise whenever named genotypes are repeatedly propagated and distributed, including materials maintained in germplasm repositories and clonally propagated crops.

## Results

### Genetic and epigenetic variation resolves relationships among Col-0 laboratory stocks

We first asked whether genetic and epigenetic variation accumulated during independent propagation could resolve relationships among contemporary Col-0 laboratory stocks. After quality filtering, the analysis included 75 Col-0 lines and two Col-1 individuals used as the outgroup (Dataset S1). Across these samples, we identified 1,415 single-nucleotide polymorphisms (SNPs), with an average of 44 SNPs per individual relative to the reference genome and a range of 6–73, and used them to construct a SNP-based phylogeny (SNP tree; Fig. 1; Dataset S2). Because relatively few nucleotide substitutions accumulate over the timescale of laboratory propagation(10, 15, 17), we complemented this analysis with single-site CG epimutations. These changes occur substantially more frequently than DNA mutations and, at the clock-like sites examined here within gene-body-methylated genes, are generally considered functionally neutral, making them abundant markers of recent lineage history(13, 17, 18). We identified 169,677 such epimutations and used them to construct an epimutation-based phylogeny (EPI tree; Fig. 1).

**Figure 1.**
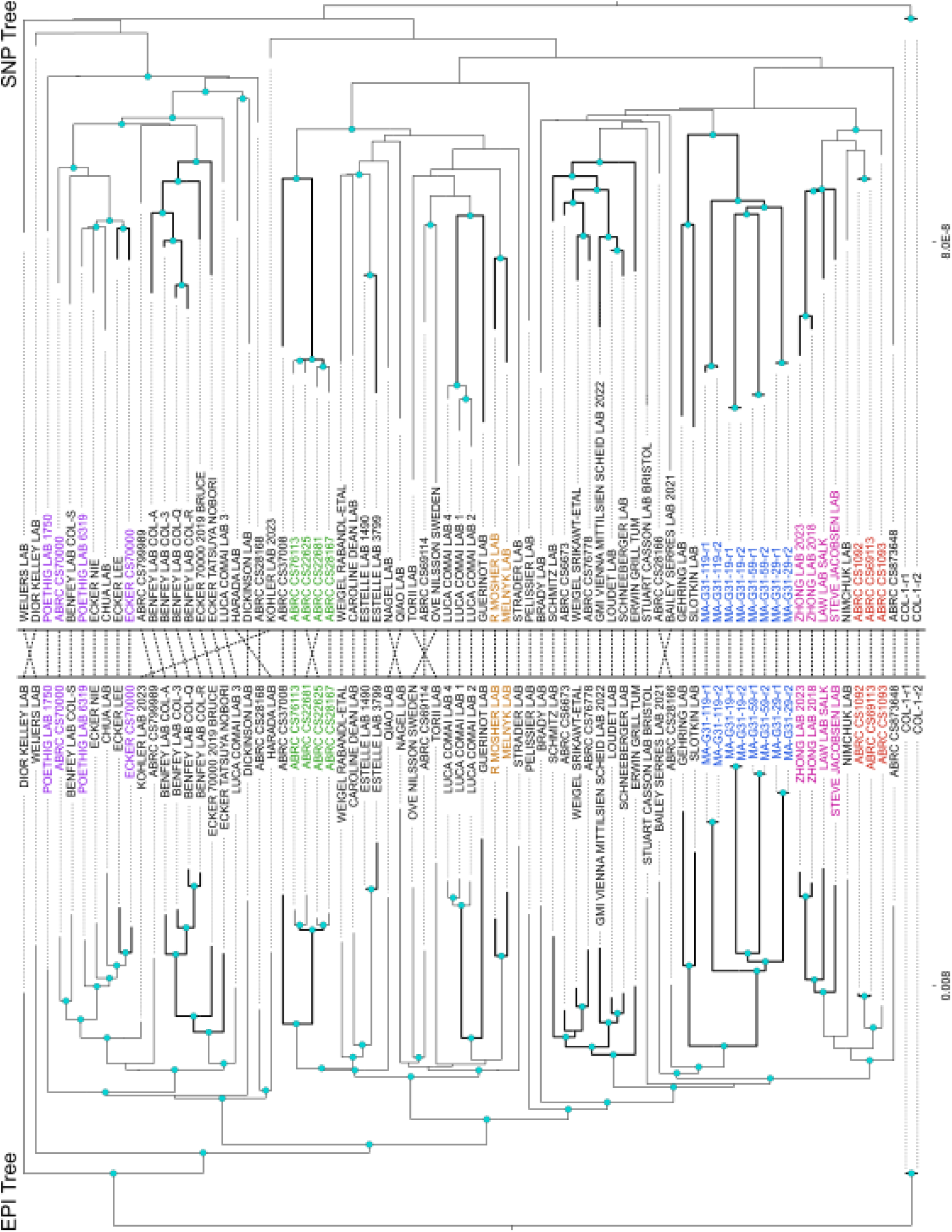
Recent divergence among Col-0 laboratory stocks. The epimutation maximum likelihood tree (EPI tree - left) was generated from 59,314 epimutations identified across 481,652 sites that exhibited clock-like properties (CpG sites within gene body methylated genes) across all individuals. The single nucleotide polymorphism maximum likelihood tree (SNP tree - right) was generated from 1,415 SNPs identified across the whole genome across all individuals. Bold branches represent relationships present in both trees with 42.1% (32/76) of the nodes in the EPI tree having the same descendant lineages in the SNP tree. The blue circles designate nodes with bootstrap support values ≥ 90. The dotted lines down the center connect matching individuals between the two trees and show that most individuals form a similar clade in each tree. Individuals are colored if they are part of a known relationship: (purple) Ecker CS70000 seeds came from Poethig, (brown) Mosher and Melnyk were both in the Baulcombe Lab (not part of dataset), (blue) mutation accumulation lines spanning 31 generations (Shaw et al., 2000), (pink) Zhong and Law were both in the Jacobsen Lab, (red) ABRC stocks CS69113 and CS1093 are direct descendants of CS1092. Branch lengths represent the number of substitutions/epimutations per site.

Despite being based on different molecular markers, the SNP and EPI trees recovered many of the same relationships, including documented ABRC pedigrees and shared laboratory histories. According to ABRC records, stocks CS28167, CS76113 and CS22681 are direct descendants of CS22625, and these four stocks formed a clade in both trees (Fig. 1, green). Stocks from Zhong and Law, both formerly in Steve Jacobsen’s laboratory, also formed a clade in both phylogenies (Fig. 1, pink), as did stocks from Mosher and Melnyk, who were both formerly in the Baulcombe laboratory (Fig. 1, brown). ABRC records further indicate that CS70000 seeds were donated by Ecker in 2008 and that Ecker’s stock originated from Poethig. Consistent with this history, the ABRC CS70000, ECKER CS70000 and POETHIG LAB 6319 stocks grouped together with four additional lines in both trees (Fig. 1, purple).

The collection also included four mutation-accumulation (MA) lineages that had been propagated independently by single-seed descent for 30 generations (9), with each lineage represented by a pair of sibling descendants (Fig. 1, blue). Because the relationships among these samples were defined by an experimental pedigree, they provided an independent reference for evaluating the inferred lineage structure. Both trees recovered the expected grouping of sibling pairs and the shared ancestry of the four MA lineages (Fig. 1). Thus, despite incomplete provenance information for many stocks, both phylogenies recovered relationships supported by stock-center records, shared laboratory histories and an experimentally defined pedigree. Genetic and epigenetic variation among Col-0 stocks therefore retained a clear record of their propagation history.

### Molecular dating places the Col-0 MRCA near its historical origin

Having established that SNPs and CG epimutations recover known relationships among contemporary Col-0 stocks, we next sought to estimate the time to their most recent common ancestor using molecular-clock approaches. Because the historical origin of the Columbia lineage is known, this system provides an opportunity to test whether SNP- and CG epimutation-based dating recovers a timescale consistent with the emergence and dissemination of Col-0. Following previous approaches, we used externally calibrated per-generation mutation and epimutation rates to convert branch lengths in the SNP and EPI trees into generations(10–13, 15, 17). Comparing these estimates with the known historical timeframe also allowed us to infer the average frequency of laboratory propagation since the establishment of Col-0.

The experimentally defined MA lineages provided an internal temporal reference. Both approaches estimated their divergence from the common ancestor of the MA group to be consistent with the known 31 generations (Table 1; Fig. 2B). When applied to the wider collection, the two molecular clocks placed the most recent common ancestor of the sampled Col-0 stocks approximately 55– 60 generations in the past (Table 1). Given that the Columbia lineage was established in the 1950s, this estimate implies an average propagation frequency of approximately one generation per year. Some contemporary stocks were separated by more than 100 generations of cumulative divergence, illustrating the extent of independent propagation represented in the collection.

**Figure 2.**
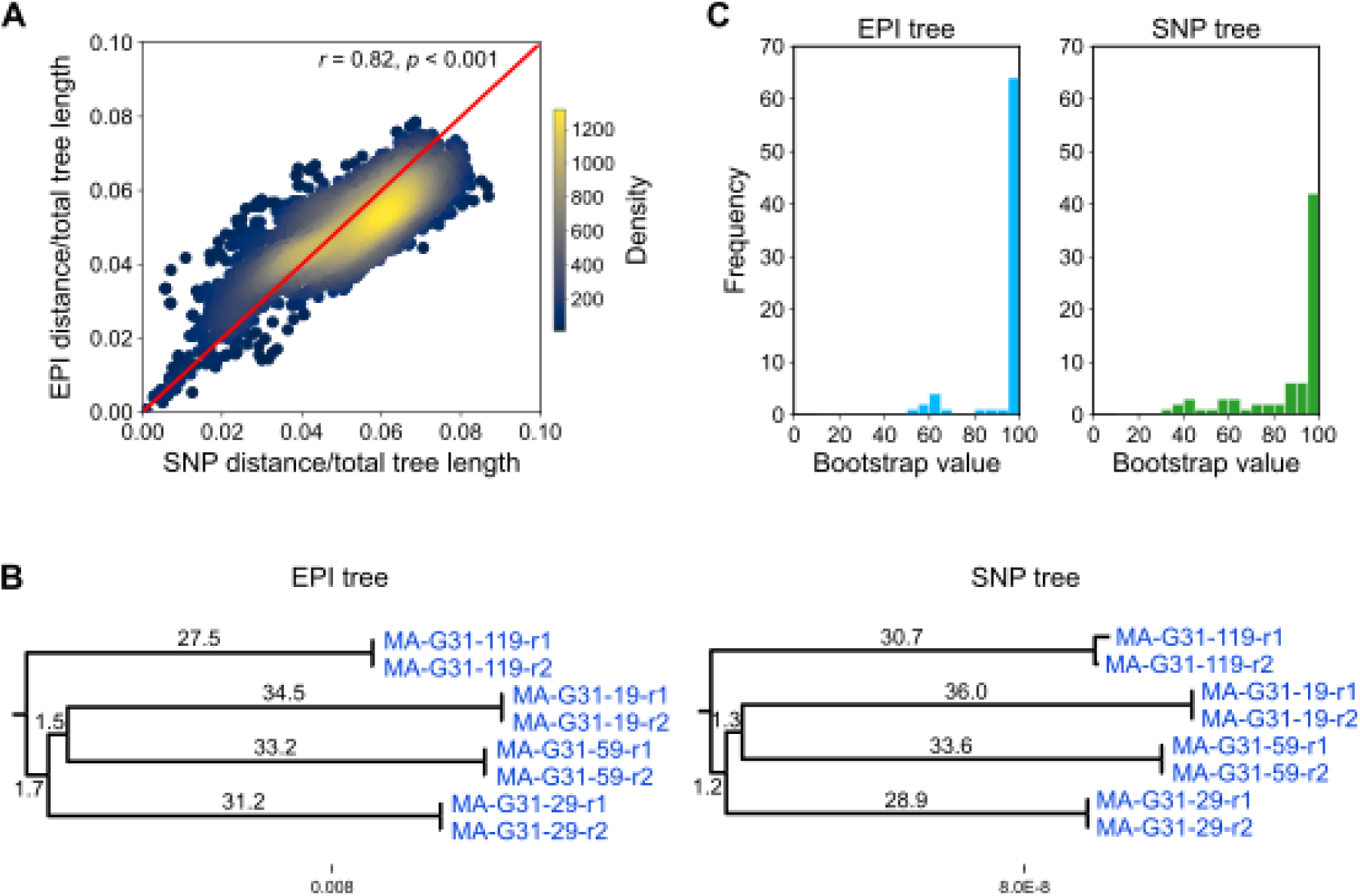
Comparison of SNP- and epimutation-based estimates of divergence. **A.** Pairwise distance comparison between EPI tree and SNP tree. Each dot represents one pair of individuals and is colored by density. The axes measure the distance (tip-to-tip) between the pair of individuals in the SNP tree (x-axis) and the EPI tree (y-axis) divided by the total tree distance for each tree respectively. Pearson’s r = 0.82, p < 0.001. **B.** Mutation accumulation (MA) line individuals from EPI tree and SNP tree with branch length values representing number of generations. **C.** Distributions of bootstrap support values for EPI tree and SNP tree. Bootstrap values were organized into bins representing intervals of five. 86.8% and 65.8% of the nodes in the EPI tree and SNP tree, respectively, have a bootstrap support value greater than or equal to 90, indicating the relationships in the EPI tree are more confident.

**Table 1.**
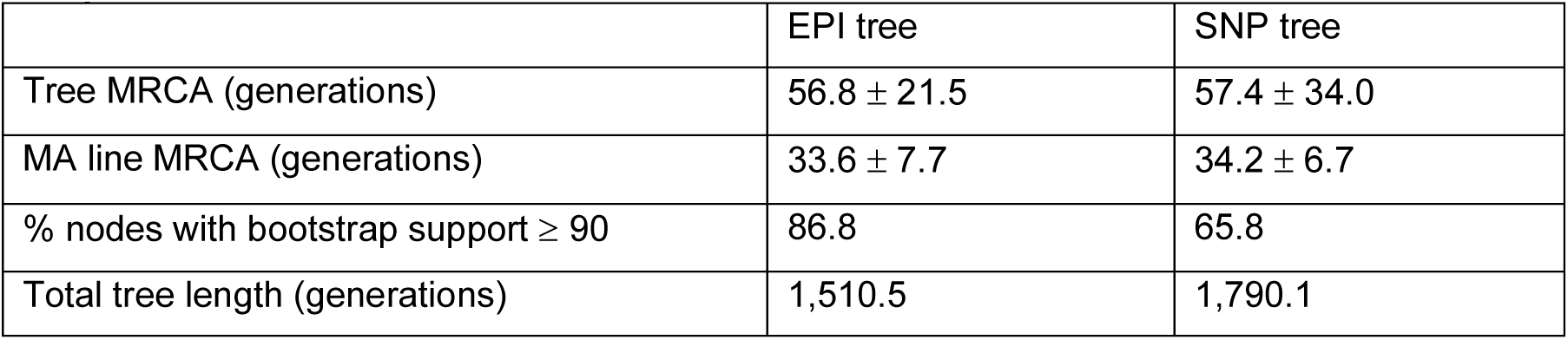
Comparison of EPI tree and SNP tree. For each tree, the estimate of the number of generations since the most recent common ancestor (MRCA) for all samples, the MRCA estimate for just the MA line clade, the proportion of nodes with bootstrap support ≥ 90, and the total tree length in number of generations. The MRCA estimates were calculated by taking the mean distance from root to tip across all samples and calculating the 95% confidence intervals.

|  | EPI tree | SNP tree |
| --- | --- | --- |
| Tree MRCA (generations) | 56.8 ± 21.5 | 57.4 ± 34.0 |
| MA line MRCA (generations) | 33.6 ± 7.7 | 34.2 ± 6.7 |
| % nodes with bootstrap support ≥ 90 | 86.8 | 65.8 |
| Total tree length (generations) | 1,510.5 | 1,790.1 |

The two marker systems also yielded similar overall divergence patterns, with strongly correlated relative pairwise distances among stocks (Pearson’s *r* = 0.82, *P* < 0.001; Fig. 2A). However, the EPI tree provided stronger statistical support: 86.8% of its nodes had bootstrap support ≥90, compared with 65.8% in the SNP tree (Fig. 2C; Table 1). This greater resolution reflects the substantially larger number of single-site CG epimutations that accumulated over the same period and supports their use as a high-resolution molecular clock over short evolutionary timescales, consistent with recent findings by Yao et al. (17).

### Characterization of accumulated genetic variation

The substantial divergence among contemporary Col-0 lineages raises the question of whether mutations accumulated during laboratory propagation are largely neutral or include variants with potential functional consequences. We therefore examined the mutation spectrum, allele-frequency distribution, genomic location and predicted consequences of the 1,415 identified SNPs. To place these patterns in context, equivalent analyses were performed using variants from the *A. thaliana* 1001 Genomes Project, which represent standing genetic variation across natural accessions and provide a broader reference for the distribution and functional constraint of SNPs (19).

We first examined whether the variants accumulated during laboratory propagation reflected the characteristic spectrum of spontaneous mutation in *A. thaliana*. The relative frequencies of the six possible base-substitution classes were broadly similar between the Col-0 collection and the 1001 Genomes dataset, with C-to-T substitutions representing the most abundant class in both (Fig. 3A). This enrichment is consistent with previous mutation-accumulation and ultra-accurate sequencing studies and is compatible with spontaneous cytosine deamination, including deamination of methylated cytosines (10, 15, 20). Thus, the mutation spectrum observed among laboratory Col-0 lineages resembles both experimentally measured spontaneous mutation and standing variation retained in natural populations.

**Figure 3.**
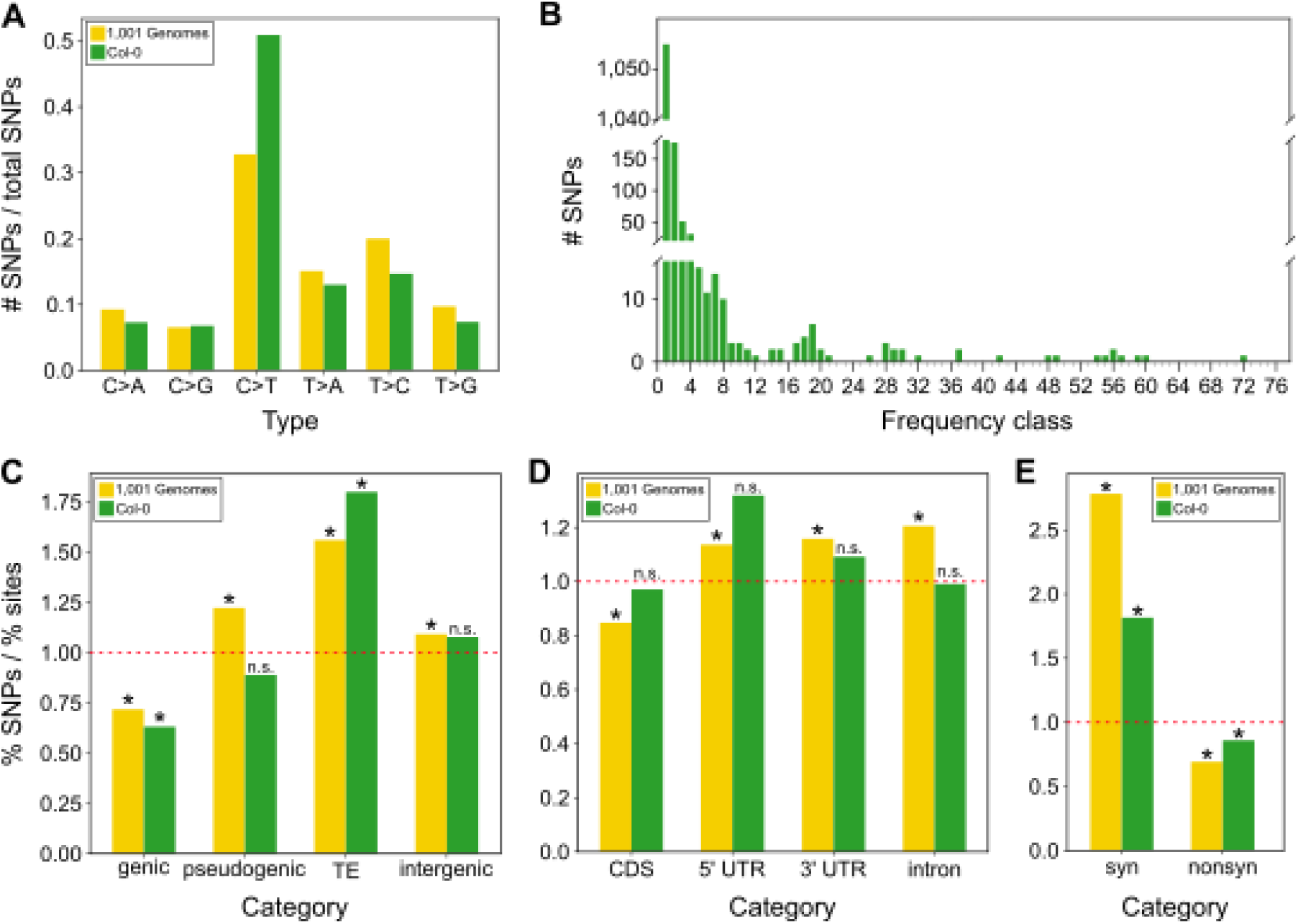
Characterization of SNPs in the Col-0 population. **A.** Site frequency spectrum showing the number of SNPs per allele frequency class. (**B, C, D**). Bar charts indicating the proportion of SNPs out of the proportion of sites falling in specific genomic categories in the 1,001 Genomes dataset (yellow) and in the Col-0 dataset (green). The proportion of sites falling into each category was determined according to the TAIR10 annotation. A value of 1 is indicated by the dotted red lines and defines the expected ratio if the SNPs were uniformly distributed. Significance was calculated using a binomial test with a Benjamini-Hochberg correction for multiple testing. An asterisk indicates a significant enrichment or depletion of SNPs in that region and “n.s” designates no significance. The 1,001 Genomes dataset contains 11,458,975 SNPs and the Col-0 dataset contains 1,415 SNPs. **E.** Mutation spectrum for Col-0 dataset (green) and 1,001 Genomes dataset (yellow) showing mostly C > T mutations.

The site-frequency spectrum was strongly skewed toward rare variants, with 1,055 of the 1,415 SNPs occurring in only a single line (Fig. 3B). This predominance of lineage-specific variants is consistent with their recent accumulation during independent propagation. To determine where these mutations occurred, SNPs were classified according to their location in genic, pseudogenic, transposable-element (TE) and intergenic regions and compared with the genomic representation of each feature class (Fig. 3C). Genic SNPs were further assigned to coding sequences, 5′ and 3′ untranslated regions, or introns, and coding SNPs were classified as synonymous or nonsynonymous substitutions (Fig. 3D–E).

In both the Col-0 and 1001 Genomes datasets, SNPs were significantly enriched in TEs and for synonymous substitutions, whereas genic and nonsynonymous SNPs were depleted relative to their genomic representation (binomial tests with Benjamini–Hochberg correction). These shared patterns suggest that the variants retained among laboratory Col-0 lineages are subject to constraints resembling those observed in natural populations, including mutation bias and possibly purifying selection against mutations with deleterious effects on gene function (20, 21).

Despite this overall depletion, the Col-0 collection contained 170 nonsynonymous substitutions and several variants predicted to have more severe consequences. SnpEff (22) classified 16 of the 1,415 SNPs as high-impact variants, including losses of start codons, gains of stop codons and splice-donor or splice-acceptor changes (Dataset S3). These variants represent candidates for functional differences among laboratory lineages, which we examine further using gene-expression data below.

### Characterization of accumulated epigenetic variation

The single-site CG epimutations used for lineage reconstruction represent only one form of heritable methylation variation. These changes arise rapidly, particularly within gene-body-methylated genes (11–13, 17, 18), and are generally considered functionally neutral (17, 23). Regional clusters of CG changes within such genes likewise appear to accumulate neutrally (24), and are not usually associated with altered transcription (23). By contrast, a rarer class of regional methylation change involves coordinated differences in CG, CHG and CHH methylation, resembling the transposable-element-like methylation (teM) typically associated with transcriptional silencing (25–30). When such changes overlap genes or regulatory regions, they may affect gene expression and potentially contribute to phenotypic variation (31). Here, we refer to these regional changes as teM differentially methylated regions (teM DMRs) or candidate epialleles.

Across 147 Col-0 individuals, we identified 1,116 teM DMRs (Dataset S4) showing differential methylation in all three sequence contexts. Of these, 202 overlapped gene bodies and were therefore classified as differentially methylated genes (Fig. 4A; Dataset S5). Because stable teM variation may be restricted to particular genomic regions, we next asked whether the number of affected genes continued to increase as additional individuals were sampled or instead approached saturation.

**Figure 4.**
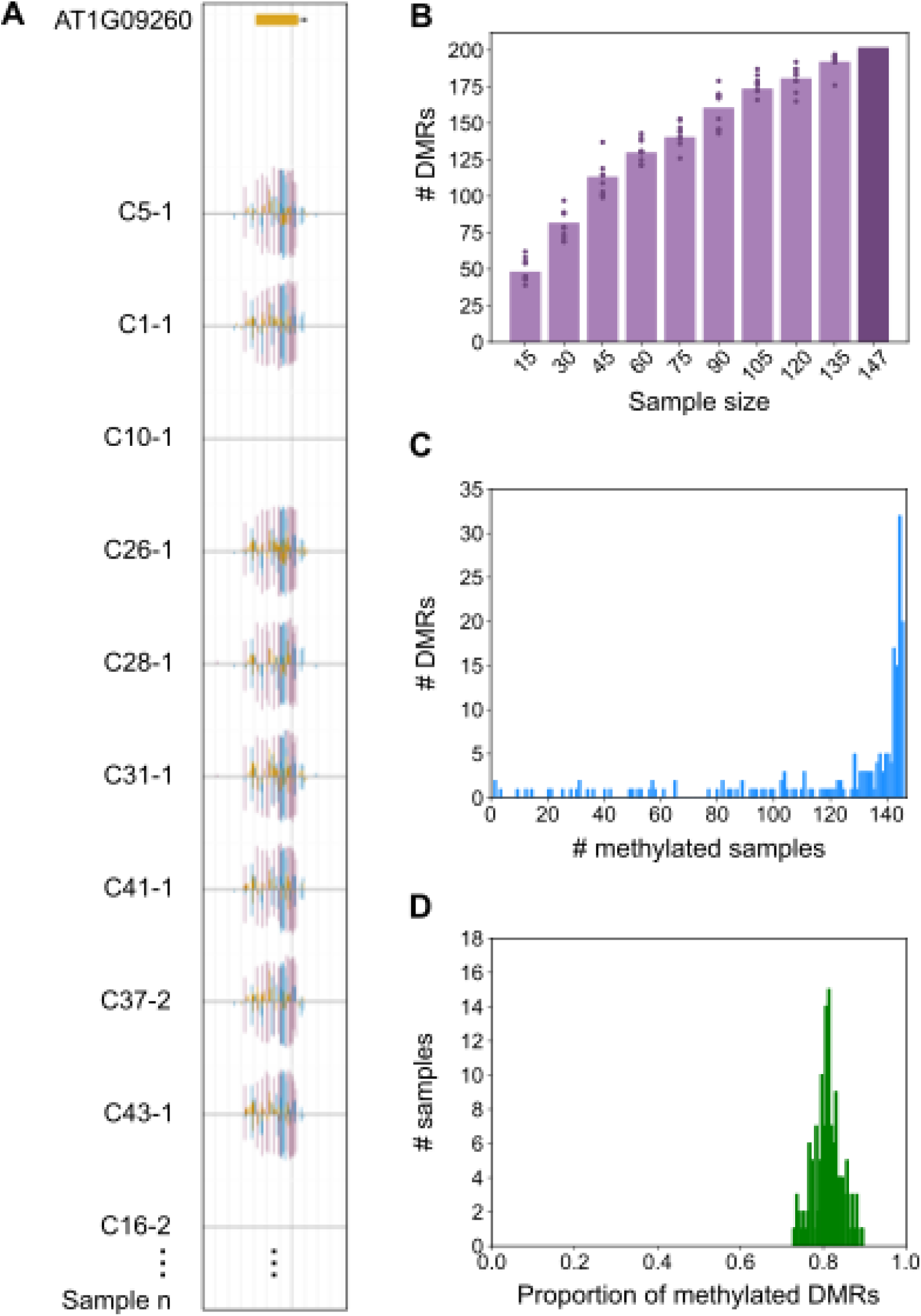
Characterization of epialleles in the Col-0 population. **A.** An example of a gene with differential methylation in all three contexts (purple: mCG, blue: mCHG, yellow: mCHH). **B.** The number of differentially methylated regions (DMRs) overlapping a gene per sample size group. Each bar, excepting the last (147), represents the mean number of DMRs found across ten randomly selected sample groups. Each dot represents the number of differentially methylated genes found in a single sample group. **C.** Methylated individual frequency spectrum showing the frequency of differentially methylated genes for which 1, 2, 3,…,146 individuals are methylated. For each differentially methylated gene, most individuals are methylated. **D.** The frequency distribution of individuals based on the proportion of differentially methylated genes for which an individual is methylated. On average, an individual is methylated at ∼80% of all identified differentially methylated genes.

The number of genes containing teM DMRs increased with sample size but began to plateau (Fig. 4B). A Michaelis–Menten model provided a close fit to the data (R² = 0.996) and estimated an asymptotic total of approximately 311 genes susceptible to teM variation in Col-0, with a sample of approximately 168 individuals projected to approach saturation. These results suggest that stable genic teM variation is largely restricted to a limited subset of genes.

We next examined the inferred direction of these regional methylation changes. Under a simple parsimony assumption, the methylation state found in the majority of individuals was treated as ancestral, such that a rare unmethylated state in an otherwise methylated region was classified as a methylation loss, and the converse as a methylation gain. Most candidate epialleles were inferred to have arisen through loss rather than gain of teM (Fig. 4C), suggesting that failure to maintain an existing methylated state is the predominant source of regional epigenetic variation among these lineages. This is consistent with a recent report of natural hypomethylated epialleles in natural accessions of *A. thaliana* being more abundant (32).

Finally, we asked whether particular Col-0 individuals showed consistently elevated or reduced methylation across candidate epialleles. No clear outlier lineages were detected. On average, an individual carried the methylated state at 81.1% of candidate epialleles, with values ranging from 72.8% to 89.6% (Fig. 4D). Thus, regional teM variation was distributed across the collection rather than being concentrated in a small number of unusually methylated or unmethylated lineages.

### Transcriptional consequences of accumulated genetic and epigenetic variation

The preceding analyses identified nonsynonymous and predicted high-impact SNPs, as well as regional teM variants with the potential to influence gene regulation. To test whether this accumulated genetic and epigenetic variation was accompanied by transcriptional differences, we generated RNA-seq data for 17 Col-0 lines selected to represent the major clades of the SNP- and epimutation-based phylogenies (i.e. the breadth of phylogenetic diversity in the collection) (Fig. 5A; Fig. S1; Dataset S6). The sampling also included two Col-1 individuals as an outgroup and, where possible, lines carrying contrasting alleles at candidate SNPs or contrasting methylation states at candidate teM DMRs. This design allowed us to examine both broad transcriptomic differentiation among lineages and expression differences associated with specific genetic or epigenetic variants.

**Figure 5.**
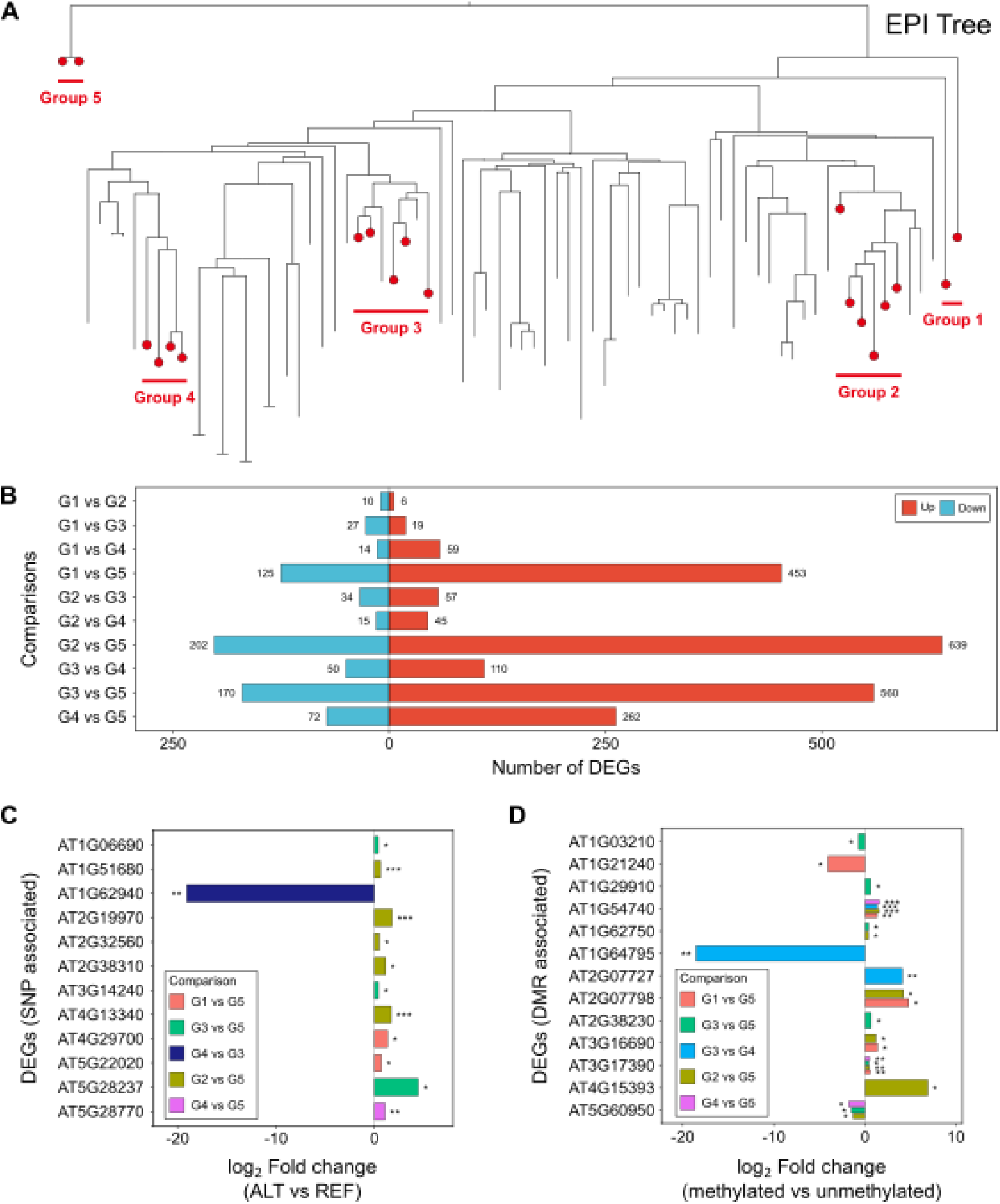
Transcriptional consequences of accumulated genetic and epigenetic variation among Columbia laboratory lineages. **A.** Tanglegram of the epimutation-based phylogeny, with the five lineage groups used for RNA- seq shown in red. Groups 1–4 correspond to Col-0 clades recovered in both trees; Group 5 comprises the Col-1 individuals used as an outgroup. **B.** Number of differentially expressed genes (DEGs) for each pairwise comparison between lineage groups, split by direction of change relative to the first group in each comparison (up, red; down, blue). DEGs were identified with DESeq2 using two-sided Wald tests and a Benjamini–Hochberg-adjusted P < 0.05. **C.** Log₂ fold changes for 12 differentially expressed genes (DEGs) associated with candidate SNPs. For each locus, fold change was calculated as the expression in lines carrying the alternative allele (ALT) relative to lines carrying the reference allele (REF). **D.** Log₂ fold changes for 13 DEGs associated with candidate transposable-element-like methylation differentially methylated regions (teM DMRs). For each locus, fold change was calculated as the expression in lines carrying the methylated state relative to lines carrying the unmethylated state. In both panels, comparison labels list the alternative-allele (C) or methylated (D) group first, so that the sign of each bar follows the order shown; colors denote the group comparison, and asterisks indicate Benjamini– Hochberg-adjusted P values (*P < 0.05, **P < 0.01, ***P < 0.001).

Pairwise differential-expression analyses between phylogenetic clades identified 16–841 differentially expressed genes (DEGs; two-sided Wald tests, Benjamini–Hochberg-adjusted P < 0.05; Fig. 5B), although the number varied substantially among comparisons. The largest numbers of DEGs occurred in comparisons involving the Col-1 outgroup, consistent with its deeper genetic and epigenetic separation from the Col-0 laboratory lineages. Comparisons involving the Col-1 outgroup identified 334-841 DEGs, with the largest number detected in Group 2 versus Group 5 (841 DEGs). Comparisons among Col-0 clades generally yielded fewer DEGs, but several still showed substantial transcriptional differentiation. Comparisons among the four Col-0 groups yielded 16-160 DEGs. The largest ingroup contrast was Group 3 versus Group 4, which contained 160 DEGs. Thus, independently propagated Col-0 stocks have accumulated measurable differences in gene expression, although these differences are smaller than those separating Col- 0 from Col-1.

We next asked whether transcriptomic divergence increased progressively with lineage divergence. For each pair of Col-0 lines or clades, the number of DEGs as an expression-divergence metric was compared with the corresponding genetic and epigenetic phylogenetic distances. Col-1 was excluded because its substantially greater divergence could dominate the relationship. No clear association was detected between transcriptomic and phylogenetic divergence among the Col-0 lineages. Neither measure of phylogenetic divergence was associated with the number of DEGs among the Col-0 groups (Pearson’s r = −0.0015, P = 1.00 for average distance; r = 0.50, P = 0.31 for MRCA distance; Fig. S3). Gene-expression differences therefore do not appear to accumulate uniformly as a simple function of propagation time. Instead, transcriptional divergence is unevenly distributed among lineages, consistent with effects concentrated at particular genetic variants, regional epimutations, or lineage-specific propagation histories.

To identify specific accumulated variants associated with expression differences, we intersected DEGs with genes carrying nonsynonymous or other potentially functional SNPs and with genes overlapping teM DMRs. We focused on variants that were shared among members of a clade but differed between clades, thereby identifying lineage-associated genetic and epigenetic states. This analysis identified 12 DEGs associated with candidate SNPs and 13 DEGs overlapping candidate epialleles (Dataset S7, S8). At these candidate SNP loci, lines carrying the alternative allele differed in mean expression from lines carrying the reference allele (Fig. 5C). Similarly, methylated and unmethylated epigenotype groups differed in expression at the genes associated with genic teM DMRs (Fig. 5D). At representative loci, genome-browser profiles showed that coordinated differences in CG, CHG, and CHH methylation across the DMR coincided with differences in RNA abundance between epigenotype groups (Fig. S4).

Among these candidates, *MTO3/SAMS3* (AT3G17390) encodes an S-adenosylmethionine synthetase, whereas *PYL4* (AT2G38310) belongs to the PYR/PYL/RCAR family of abscisic acid receptors. These associations do not establish that the SNPs or methylation variants directly caused the observed expression differences, because other linked lineage-specific changes may contribute. Nevertheless, they identify specific loci at which accumulated genetic or epigenetic variation is associated with transcriptional consequences. Independently propagated Col-0 lineages therefore differ not only in genome sequence and DNA methylation, but also at a subset of transcriptionally variable loci that could contribute to phenotypic differences among laboratory stocks.

## Discussion

Our findings challenge the common assumption that Col-0 represents a single, invariant reference genotype. Decades of seed exchange and independent propagation have instead produced a network of related laboratory lineages that retain the same accession name while differing genetically, epigenetically and transcriptionally. This distinction matters because experiments performed in nominally identical Col-0 backgrounds may not always begin from the same molecular state.

The genetic variation observed among these lineages has several features expected of recent spontaneous mutation, but its distribution also indicates that laboratory propagation is not evolutionarily neutral. The depletion of genic and nonsynonymous variants is consistent with mutation bias (20, 21) and possibly purifying selection, whether through reduced viability, fertility or inadvertent selection during routine propagation. At the same time, potentially consequential mutations can persist within particular stocks. This provides a plausible explanation for previous reports in which phenotypic differences among Col-0 isolates could be traced to fixed background variants (16, 33). The practical implication is not that all Col-0 stocks are functionally distinct, but that lineage-specific variants should be considered when phenotypes fail to reproduce across laboratories.

The transcriptomic analysis supports this more nuanced interpretation. Gene-expression differences did not increase uniformly with phylogenetic distance, suggesting that functional divergence is not a simple cumulative consequence of propagation time. Rather, transcriptional differences appear to be concentrated at particular loci and in particular lineages. Associations between expression, candidate SNPs and regional methylation variants identify plausible molecular contributors, but they do not by themselves establish causality because other linked changes may underlie the observed differences. Functional validation will therefore be required to distinguish causal variants from lineage markers.

The epigenetic analyses also highlight the importance of distinguishing different forms of methylation variation. Single-site CG epimutations accumulated rapidly and provided a dense record of recent propagation history, but they are generally expected to be functionally neutral at the sites examined here. By contrast, regional teM variants were much rarer and restricted largely to a limited set of susceptible loci. Their predominance as methylation-loss events suggests that erosion of existing methylated states is more common than the establishment of new genic teM states. Whether this asymmetry reflects differences in underlying mutation rates, selective constraint or both remains unresolved. The apparent saturation of genic teM variation further suggests that stable epigenetic divergence may be constrained by locus-specific chromatin properties rather than occurring uniformly across the genome.

The concordance between SNP- and CG epimutation-based phylogenies shows that both marker systems retain a record of laboratory propagation. The greater resolution of the epimutation-based tree is consistent with the higher rate at which single-site CG methylation changes accumulate and supports their use for reconstructing recent lineage histories (17, 18, 34, 35). However, the inferred dates should not be interpreted as exact. Propagation intervals differ among laboratories, stocks may spend variable periods in storage, and molecular-rate estimates introduce additional uncertainty. The agreement with the known history of the Columbia lineage is therefore best viewed as evidence that the inferred timescale is broadly realistic.

Several sources of variation remain incompletely captured. Short-read sequencing is poorly suited to comprehensive detection of structural variants, which may influence both phenotype and DNA methylation. The regional methylation analysis focused on gene-overlapping teM variants, whereas non-coding DMRs may also alter regulatory activity. In addition, RNA-seq was performed on a subset of stocks and under a single experimental setting, so lineage-specific expression differences may depend on tissue, developmental stage or environment. The present results therefore likely underestimate the full extent of functional divergence among Col-0 stocks.

Similar problems have been documented in laboratory lineages of *Caenorhabditis elegans*, *Drosophila* cell lines and human cell cultures, where accumulated variation can alter phenotype or experimental response (36–40). Our results extend this concern to a widely used plant reference genotype. Careful recording of stock provenance, use of appropriately matched controls and periodic replacement from well-documented source material may therefore improve reproducibility across laboratories.

The same issue extends beyond experimental model systems. Germplasm repositories and clonal propagation programs also maintain named genotypes over long periods, often under the assumption that identity is preserved through propagation. Although such systems preserve genealogical continuity, they do not necessarily preserve complete molecular identity. Recognizing maintained genotypes as evolving lineages rather than static materials may therefore be important both for experimental reproducibility and for the long-term management of biological collections.

## Materials and Methods

### Plant Material and DNA Isolation

A sample of 78 *A. thaliana* Col-0 stocks were used in this study (Dataset S1). These Col-0 stocks include 54 seed stocks from 39 different labs (some stocks came from the same lab), 16 different Col-0 accessions ordered from the ABRC, and 8 sibling pairs representing 4 lines of the 30- generation MA lines propagated by Shaw et al. (9). The MA line individuals were included as a validation for the rest of the tree as their pedigree is known. Additionally, we used the Col-1 accession as the outgroup, as Col-0 is reported to be a direct descendant of Col-1 (41). We included two Col-1 siblings increasing our total Columbia sample size to 80. For each stock, plants were grown in Sungro soil with Osmocote fertilizer under 16 hours of light and propagated for one generation. For two progenies of each plant (siblings), DNA was isolated from leaf tissue with the Qiagen DNeasy Plant Mini Kit according to manufacturer’s instructions.

### Library Prep and Sequencing

Whole genome bisulfite sequencing (WGBS) libraries were prepared following the MethylC-Seq protocol. The DNA was sonicated to 200bp by the Georgia Genomics and Bioinformatics Core using a Covaris E220 Evolution focused ultrasonicator. The fragmented DNA was end-repaired with the Epicentre End-It DNA end-repair kit. A-tailing was done with NEB Klenow 3’-5’ exo- enzyme and adapters were ligated using NEB T4 DNA Ligase. After adapter ligation, the DNA was bisulfite converted using the EZ DNA Methylation-Gold kit and amplified using the Roche KAPA HiFi Uracil + Readymix Polymerase. Paired-end 150 reads were sequenced using the Illumina Nova-seq X plus to an average read depth of ∼20X.

To prepare the whole genome sequencing (WGS) libraries, the DNA was fragmented using a Tn5 enzyme pre-loaded with universal adapters and 5X TAPS tagmentation buffer. For 3.5 uL of each DNA sample, 0.083 uL Tn5 enzyme, 0.42 uL 5X TAPS buffer, and 0.997 uL dH2O was added. The Tn5 reaction was run in a thermocycler for 15 minutes at 55°C. The fragmented DNA was then PCR amplified using 12.5 uL 2X Taq HF Master Mix, 6.95 ul dH2O, 0.05 uL 100uM i5 indexed adapters, and 0.5 uL 10uM i7 indexed adapters. The PCR cycles were 1x 72°C (3m), 1x 95°C (1 min), 18x (95°C (10s), 55°C (20s), 72°C (3m)), 1x 72°C (5m). The libraries were then pooled, and bead cleanup was done with SPRI beads. Paired-end 150 reads were sequenced using the Illumina Nova-seq X plus to an average read depth of ∼36X.

### SNP Identification

We used FastQC v0.11.9 (42) for quality-assessment of the raw sequenced paired-end WGS reads. To identify SNPs across this Columbia WGS dataset, we used Fastp v0.23.4 (43) to trim adapter sequences and required a minimum read length of 36 bp. The paired-end reads were aligned to the TAIR10 *A. thaliana* reference genome using Bowtie2 v2.5.2 (44) (Dataset S9). Due to a high duplication rate, we merged the reads of each sibling pair to represent one individual from each seed stock. We then downsampled reads so that each individual had a maximum read depth of 30 (except for ABRC CS22681, the MA lines, and the Col-1 individuals), resulting in an average read depth of 26.1 across all individuals. We also set a minimum average read depth of 10 and removed three individuals that did not meet this threshold, reducing the sample size from 80 Columbia stocks to 77. We used SAMtools v1.17 (45) to remove duplicate reads, keeping only properly paired and uniquely aligned reads. Moreover, reads with a MAPQ score less than 30 were removed. We used BCFtools v1.15.1 (45) mpileup (-Q 30) and call commands to generate a VCF file for each individual, retaining only homozygous SNPs that had a minimum mapping quality of 30, a minimum quality score (VCF QUAL) of 30, and a minimum read depth of 5. Then we merged the individual VCF files into a single VCF file and removed any multiallelic or monomorphic sites (bcftools view -m2 -M2). Finally, we retained only SNPs at positions where more than half the individuals had a base call and removed positions for which the alternative allele was called for all individuals (Dataset S2).

### SNP-based Phylogeny

To generate a maximum likelihood phylogenetic tree from SNPs, we first used BCFtools v1.15.1 (45) to generate a consensus sequence for each individual. The consensus sequence for each individual was the TAIR10 reference genome with the alternative allele replacing the reference allele at each SNP position. The consensus sequences were concatenated into a multiple sequence alignment and used as input for generating a maximum likelihood tree using the GTR+G12 substitution model in IQ-TREE2 v2.2.2.7 (46) with 1,000 bootstrap replicates (arguments: -m GTR+G12 -nt AUTO -B 1000). GTR+G12 argument specifies the general time reversible substitution model and a discrete Gamma rate heterogeneity across sites model with 12 rate categories. The bootstrap support values were generated using the Ultrafast Bootstrap approximation. The tree was visualized in FigTree with the two Col-1 individuals chosen as the root. The branch lengths were scaled by a factor of 1/μ where μ = 7 × 10⁻⁹ (10).

### Epimutation Identification

We used FastQC v0.11.9 (42) for quality-assessment of the raw sequenced paired-end WGBS reads. We used Methylpy v1.4.6 (47) to align the reads to the TAIR10 *A. thaliana* reference genome and produce methylation calls for every cytosine in the genome. Methylpy used Cutadapt v2.8 (48) to trim adapters and Bowtie2 v2.5.2 (44) to perform the alignment. The *A. thaliana* chloroplast was used as the unmethylated control to calculate the non-conversion rate (Dataset S10). We processed the Methylpy allc output files for each line with Methimpute v1.34.0 (49) to generate posterior probabilities for the methylation status of each cytosine. We merged the Methimpute output files into three new files: (1) a coverage matrix, (2) a methylation status matrix, and (3) a posterior probability matrix (collect_methimpute_results_largefiles.py). Each of these three matrices was reduced to only clock-like regions by retaining only CG sites found in Col-0 gbM genes totaling 481,653 sites (annotation.py). The list of Col-0 gbM genes was previously defined by Zhang et al. (30). We then filtered the methylation status matrix based on coverage and posterior probability (qualification.py). Methylation calls covered by at least one read and with a posterior probability greater than or equal to 0.8 were retained (methylated represented as “1”, unmethylated represented as “0”). The methylation calls for cytosines not meeting these qualifications were replaced with “?”. The final filtered matrix (*-hetero-masked.tsv) had rows defined by cytosine position and columns defined by Columbia stock. Each cell had either “0”, “1”, or “?” designating homozygous unmethylated, homozygous methylated, or missing, respectively.

### Epimutation-based Phylogeny

We generated two epimutation-based phylogenetic trees; one with sibling pairs for each Columbia stock (Fig. S1) and one with a sample composition to match the snp-based phylogenetic tree (retaining only one sibling from each Col-0 pair, excepting the MA line siblings, and removing the same three previously removed stocks). To generate each of these trees, the methylation status matrix was converted to a single binary FASTA multiple sequence alignment representing each Columbia stock (one alignment per sample set) with “1” indicating methylated, “0” indicating unmethylated, and “?” indicating a missing call (df_to_fasta.py). This alignment was then used as input for generating a maximum likelihood tree using the GTR2+G12 substitution model in IQ- TREE2 v2.2.2.7 (46) with 1,000 bootstrap replicates (arguments: -m GTR2+G12 -nt AUTO -B 1000). GTR2+G12 specifies the general time reversible substitution model for binary data and a discrete Gamma rate heterogeneity across sites model with 12 rate categories. The bootstrap support values were generated using the Ultrafast Bootstrap approximation. The resulting tree was visualized in FigTree with the two Col-1 individuals chosen as the root. The branch lengths were scaled by a factor of 1/μ where μ = 1.2 × 10⁻³ (50). This epimutation rate estimate includes epi- heterozygous sites, which have higher epimutation rates, whereas the previously reported estimate (∼4 × 10⁻⁴) excluded these sites. The average number of epi-homozygous methylated sites, epi-homozygous unmethylated sites, and sites with missing methylation calls per individual was 108,687, 272,814, and 100,151, respectively (Dataset S11).

### Tree comparisons

To facilitate direct comparison between the SNP- and epimutation-based phylogenies, branch lengths were converted from substitutions or epimutations per site to generations by scaling with the mutation rate: 7 × 10⁻⁹ substitutions per site per generation (10), or epimutation rate: 1.2 × 10⁻³ epimutations per site generation (50), respectively. For each tree, we used the R package adephylo (51) to calculate the root-to-tip distances (Dataset S12) and the R package ape (52) to calculate the total tree lengths (Table 1) and pairwise tip-to-tip distances (Fig. 2A). Because the total tree distance of each tree differed by ∼300 generations, we normalized the pairwise distances of each tree by the respective total tree length. To assess how well the pairwise distances correlated between the two trees, we plotted for each pair the relative distance in the SNP-based phylogeny on the x-axis and the relative distance in the epimutation-based phylogeny on the y-axis. Finally, to evaluate the similarity and support of the two topologies, we calculated the proportion of nodes in each tree that had the same descendant lineages and the proportion of nodes with a minimum bootstrap support value of 90.

In both trees, the MA line individuals (Fig. 1, blue) are the only individuals for which divergence times are known. Each individual is 31 generations diverged from the most recent common ancestor (MRCA) of the MA line group. For each tree, we calculated the time of the MRCA of the MA line clade by taking the mean across each root to tip distance (Fig. 2B) and calculating the 95% confidence interval. We then performed the same calculation across all individuals for each tree to estimate the time of the MRCA of the whole group.

### DMR Identification

We used jDMRgrid v0.2.4 (53) to identify differentially methylated regions (DMRs) across all Col-0 individuals. We defined 100 bp non-overlapping windows across the whole genome and required a minimum of 10 differentially methylated cytosines per window. For each context, jDMRgrid outputs a state call matrix with a binary call (methylated/unmethylated) for each window for each individual and a posterior probability matrix with the posterior probabilities associated with each methylation call. Using the state call matrix, we removed windows showing no differential methylation. Then we used Bedtools v2.31.1 (54) to find the intersection of all three contexts, retaining only windows that showed differential CG, CHG, and CHH methylation. Defining the methylation level for each window as the number of methylated reads out of the total number of reads based on the allc files for each individual, we removed windows for which the difference in methylation level between the most and least methylated individuals was less than 30%. Additionally, for each window, we required the most and least methylated individuals to have a minimum posterior probability of 0.95. Finally, windows within 500 bp were merged.

To determine if this dataset captures all possible epialleles, we identified DMRs that overlap gene bodies for groups of increasing sample size to test for saturation. From the whole dataset of 147 Col-0 individuals, subsample groups were generated by randomly selecting 15,30,…,135 individuals. For each subsample size, 10 randomly selected groups were generated. For each subsample group, we subset the jDMRgrid matrices to only include the individuals in that group and filtered using the same steps previously described. Since the jDMRgrid binary state calls were generated per 100bp window per context, we chose the first 100 bp window from the CG context for each potential epiallele as the representative state calls for that region.

### RNA-seq analysis

RNA-seq libraries were sequenced on an Illumina platform to generate 150-bp paired-end reads. Raw reads were quality-assessed using FastQC v0.11.9 (42) and trimmed using Trimmomatic v0.39 (55) to remove sequencing adapters and low-quality bases. Trimmed paired-end reads were aligned to the TAIR10 reference genome using STAR v2.7.6a (56). Alignments were sorted and indexed using SAMtools v1.17 (45). Gene-level read counts were generated using HTSeq-count v0.13.5 (57) with the Araport11 annotation.

Count matrices were restricted to nuclear genes on chromosomes 1–5 and analyzed using DESeq2 v1.30.1 (58). For visualization purposes, raw gene counts were transformed using the regularized logarithm (rlog) method. To account for technical confounding factors, the first two principal components (PC1 and PC2) derived from an unguided principal component analysis (PCA) of rlog-normalized counts were included as covariates in the DESeq2 design matrix (design = ∼ PC1 + PC2 + tree_group). Pairwise differential expression comparisons were performed among the five phylogenetically defined clades (Group 1 to Group 5). Pairwise differential-expression analyses were performed among all five lineage groups using two-sided Wald tests implemented in DESeq2. Differentially expressed genes were defined using a Benjamini–Hochberg-adjusted P value below 0.05.

## Data availability

Data analysis scripts can be found at https://github.com/atadros123/Col-0-lab-lineages.

## Acknowledgments

We would like to thank all laboratories whose seed stocks were included in this study, Yangyang Xu for performing DNA and RNA extractions and preparing all libraries for sequencing, Kelly Dyer, Jim Leebens-Mack, Liang Liu, and Mark Minow for data analysis and visualization suggestions,

Cullan Meyer for sharing a list of Col-0 synonymous/nonsynonymous sites and providing feedback on the manuscript, as well as Xiang Li and Xuan Zhang for providing feedback on the manuscript, and Kevin Sun for telling jokes. The computational resources used in this study were provided by the Georgia Advanced Computing Resource Center. This research was supported by the NSF (MCB-2242696) and the University of Georgia Office of Research awarded to R.J.S. as well as the National Institute for General Medical Sciences of the NIH awarded to A.M.T (5T32GM142623-04).

## Supplementary Figures

**Fig. S1.**
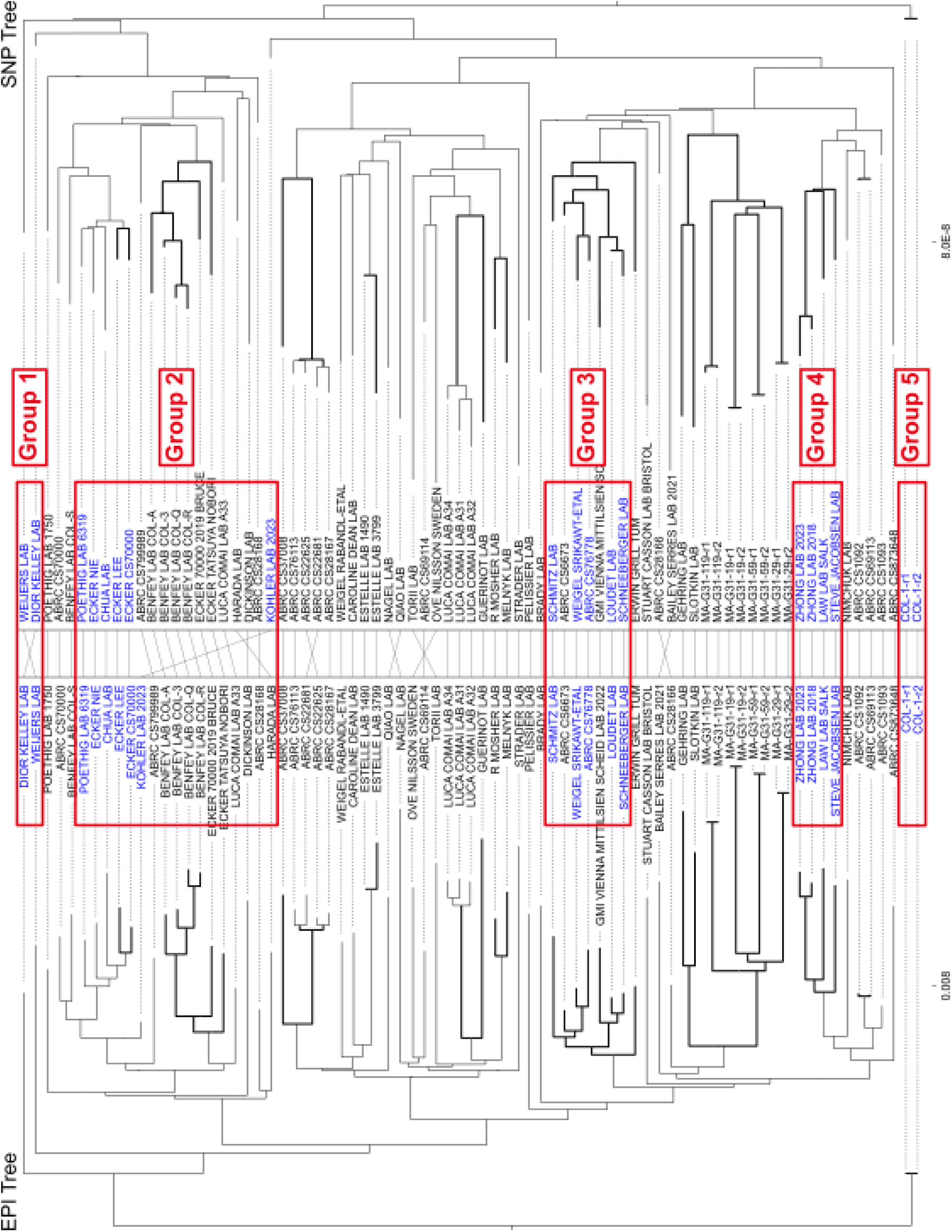
Samples selected for RNA-seq analysis. 17 Col-0 stocks and the two Col-1 replicates were selected and are colored in blue. Groups are desiganted by the red boxes.

**Fig. S2.**
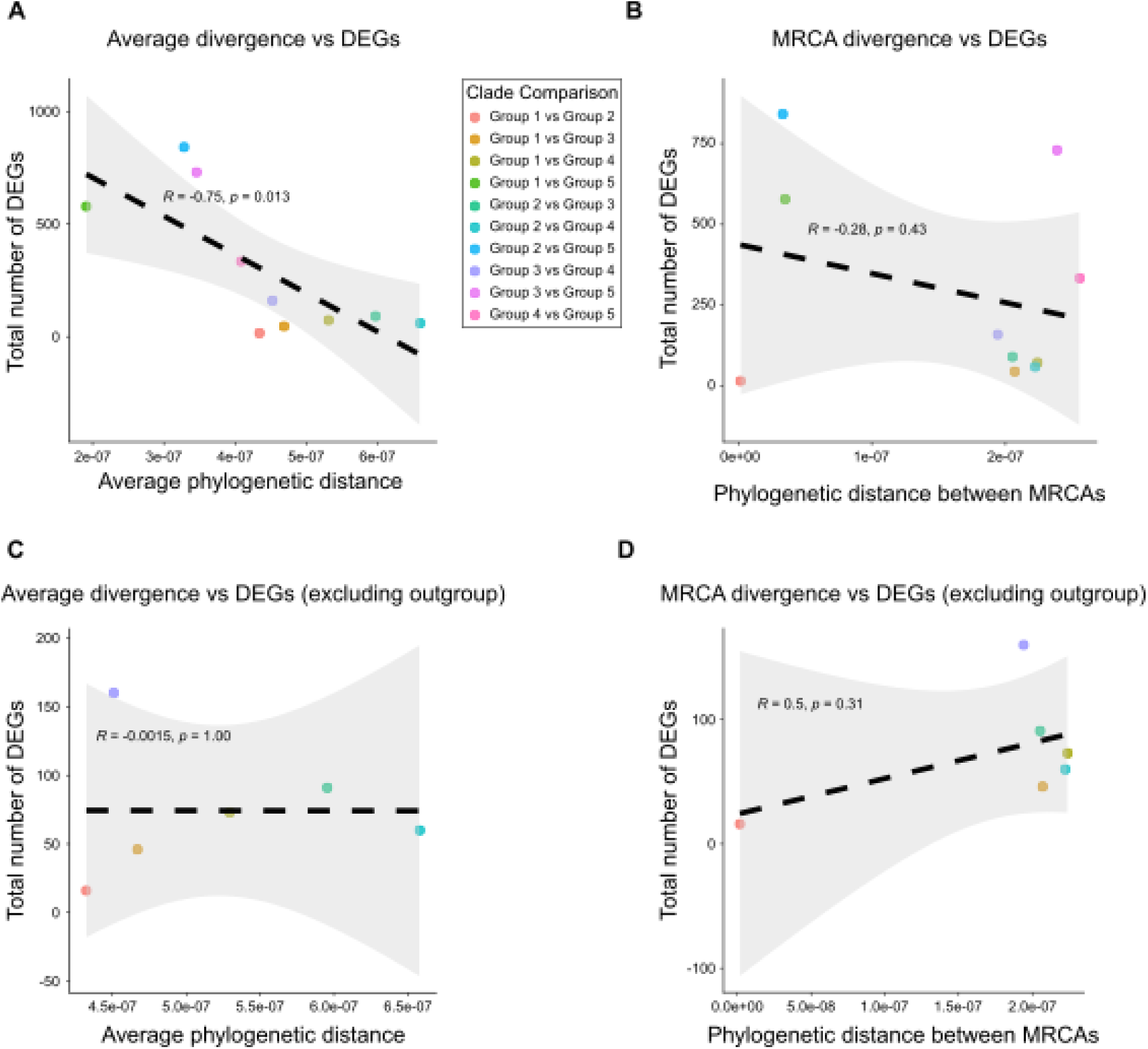
Relationship between phylogenetic divergence and transcriptional differentiation among lineage groups. The total number of differentially expressed genes (DEGs) was compared with the average phylogenetic distance between clades (A, C) or the phylogenetic distance between their most recent common ancestors (MRCAs; B, D). Analyses were performed across all pairwise clade comparisons (A, B) and after excluding comparisons involving the Col-1 outgroup (Group 5) (C, D). Each point represents one pairwise comparison. Dashed lines show linear regressions, with grey shading indicating 95% confidence intervals. Pearson correlation coefficients and corresponding two-sided P values are shown.

**Fig S3.**
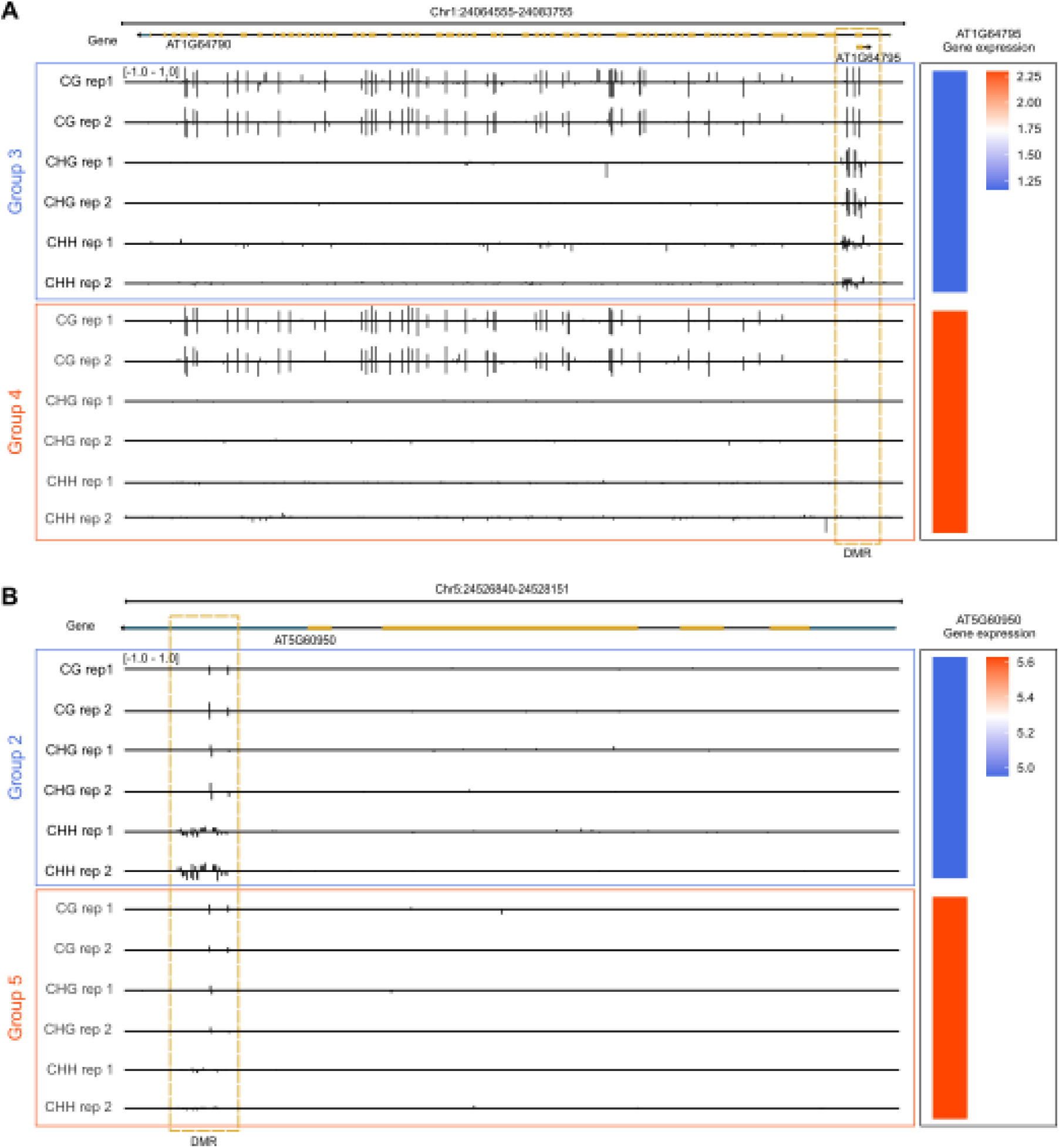
Representative candidate epialleles associated with lineage-specific gene-expression differences. A. Genome browser view of the AT1G64795 locus. Increased CG, CHG, and CHH methylation across the boxed teM differentially methylated region (DMR) in Group 3 coincided with lower expression of AT1G64795 relative to Group 4. B. Genome-browser view of AT5G60950. Increased methylation across the boxed teM DMR in Group 2 coincided with lower expression of AT5G60950 relative to Group 5, the Col-1 outgroup. WGBS-derived methylation profiles are shown for two biological replicates per group in the CG, CHG, and CHH sequence contexts. Colored bars on the right indicate the group-mean rlog-transformed expression of the indicated gene, with blue and red representing lower and higher expression, respectively.

**Fig S4.**
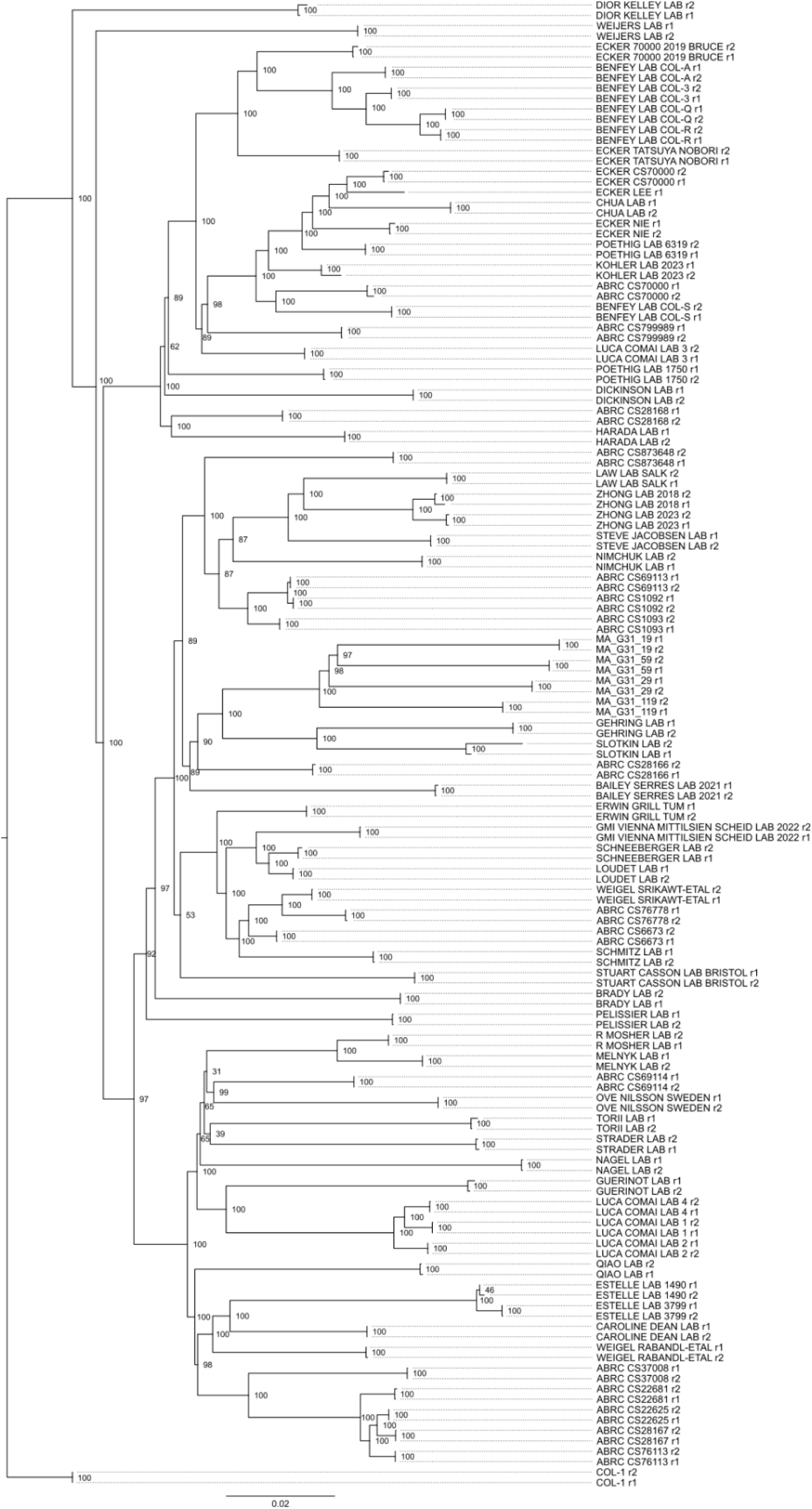
Epimutation-based phylogenetic tree including replicates. Replicates of all 78 Col-0 stocks as well as the MA lines and Col-1 outgroup. Bootstrap support values are shown at the nodes.

